# Quantifying biochemical reaction rates from static population variability within complex networks

**DOI:** 10.1101/2021.08.30.458258

**Authors:** Timon Wittenstein, Nava Leibovich, Andreas Hilfinger

**Affiliations:** Department of Physics, University of Toronto, 60 St. George St., Ontario M5S 1A7, Canada; Department of Physics, Johannes Gutenberg University of Mainz, Staudingerweg 7, 55128 Mainz, Germany; Department of Chemical & Physical Sciences, University of Toronto, Mississauga, Ontario L5L 1C6, Canada; Department of Mathematics, University of Toronto, 40 St. George St., Toronto, Ontario M5S 2E4; Department of Cell & Systems Biology, University of Toronto, 25 Harbord St, Toronto, Ontario M5S 3G5

## Abstract

Quantifying biochemical reaction rates within complex cellular processes remains a key challenge of systems biology even as high-throughput single-cell data have become available to characterize snapshots of population variability. That is because complex systems with stochastic and non-linear interactions are difficult to analyze when not all components can be observed simultaneously and systems cannot be followed over time. Instead of using descriptive statistical models, we show that incompletely specified mechanistic models can be used to translate qualitative knowledge of interactions into reaction rate functions from covariability data between pairs of components. This promises to turn a globally intractable problem into a sequence of solvable inference problems to quantify complex interaction networks from incomplete snapshots of their stochastic fluctuations.

---

Quantifying interactions between components within a complex network from snapshots of their activity is a challenge common to many areas of science. For example, understanding cellular processes requires quantifying biochemical reaction rates between molecules while typical high-throughput methods such as single-cell sequencing [1, 2], flow cytometry [3–5], or a combination thereof [6] generate static population snapshots of a subset of cellular components.

Covariability of components within cellular processes is typically analyzed using statistical associations [7–11] because accurate mechanistic modelling of biochemical reactions is impractical for complex systems due to the large number of unknown parameters and interactions [12]. However, fluctuations in biochemical reaction networks emerge from underlying physical interactions that affect each component’s rate of production and degradation rather than its instantaneous concentration. How molecular components affect each other is then difficult to infer from statistical associations [13], especially in the absence of perturbation experiments [14, 15]. For example, even perfectly linear *rate* dependencies will lead to non-linear statistical relations between observed *values* of cellular components.

We introduce a novel data analysis approach to deduce rate dependencies one interaction at a time using incompletely specified mechanistic models, see Fig. 1. Our approach exploits a local qualitative understanding of network interactions through probability balance equations [16] that must be satisfied as long as we know how one component is made and degraded. As a proof-of-principle we analyzed four example systems with markedly different global dynamics. We show numerical evidence that in those systems we could successfully infer how one component affects the production rate of another from their observed joint probability distribution without making any assumptions about the dynamics of the non-observed components.

**FIG. 1.**
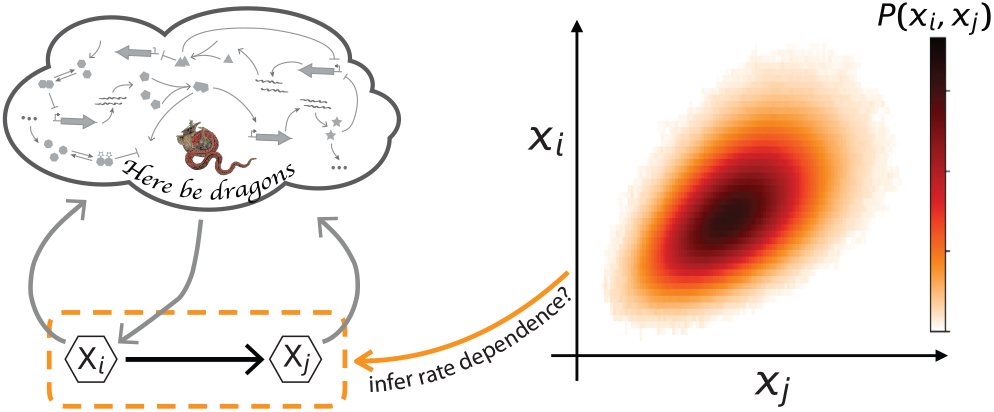
Determining how one component affects the production rate of another from observing their joint probability distribution. The joint probability distribution *P*(*x_i_, x_j_*) varies enormously between systems even for identical effects of X_*i*_ on X_*j*_. Nevertheless, all systems with a given interaction between the two components satisfy invariant probability flux balance relations [16]. Here, we demonstrate that such relations can translate empirically observed joint probability distributions into interactions between X_*i*_ and X_*j*_ even when the dynamics within the rest of the network is unknown.

## Background theory

Describing the dynamics of some components within a complex interaction network in which the interactions between many components are unknown may seem impossible. However, we can trivially do so as long as we are content with describing one component’s dynamics in terms of components directly affecting it. The actual dynamics are fundamentally indeterminable for incomplete models but if components of interest are experimentally measurable their empirically observed covariability can be used to close the problem and constrain interaction rates as described below.

We follow the previously established approach [16] to characterize “local” system dynamics within a completely general complex reaction network with probabilistic events

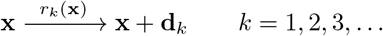

where the state vector x = (*x*_1_, *x*_2_, *x*_3_, …) of abundances can be arbitrarily high-dimensional, the *k^th^* reaction changes levels of component X_*i*_ by *δ_ki_*, and the reaction rates *r_k_* (x) are arbitrarily non-linear functions of the state vector. This notation is motivated by biochemical reaction networks but many areas of science encounter stochastic systems whose dynamics are determined by the corresponding general chemical master equation

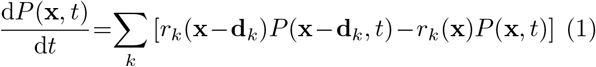

where *P*(*x, t*) denotes the probability of the system to be in state x at some time *t*. Eq. (1) is generally intractable for two reasons: first, any non-linear rate *r_k_* renders it analytically unsolvable, second, for any actual complex reaction network we never know all the rate functions.

However, even when many details of a system are unknown, any given molecular component X_*i*_ that reaches a time-independent stationary state with probability distribution *P_ss_*(*x_i_*) must satisfy

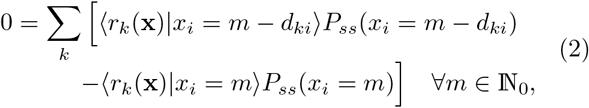

which follows from simple summation of Eq. (1) over all other variables and has been derived and discussed previously [16, 17]. In this paper, we demonstrate that Eq. (2) can be exploited to infer rate functions even when we know nothing about the dynamics of all other components X_*j*_ for *j* ≠ *i* such that conditional rates cannot be predicted from incompletely specified models.

Note, Eq. (2) is not an approximate coarse-graining but corresponds to an exact balance relation for any variable in a larger complex system at stationarity. Whether this relation applies only depends on whether the system has reached stationarity which can be verified experimentally from population snapshots taken at different timepoints. Thus the only dynamics excluded from our analysis is transient behaviour such that stationary probability distributions are not accessible from experimental data. Note, explicitly time-varying systems, such as deterministic oscillations, satisfy Eq. (2) when considering their time-averaged probability distributions and rates [16].

## Results

When the rates of all reactions directly changing X_*i*_-levels are known, Eq. (2) represents a self-consistency check that must be satisfied by the observed joint probability distribution between X_*i*_ and all variables directly affecting those rates. Next, we show how this relation can be “inverted” to determine rate functions from observed probability distributions.

While Eq. (2) must provably hold for all stationary states, any empirically observed distribution will exhibit sampling errors which can have significant effects. For example, any real experiment will have some maximum value *m*_max_ for which X_*i*_ is observed and thus Eq. (2) will clearly be violated for unbounded systems because Eq. (2) cannot balance when *m* = *m*_max_ due to the lack of sampling of the rarest states. Inverting the equation system Eq. (2) to identify the functional dependencies of *r_k_* (x) thus requires minimizing deviations from the predicted relations. In general, minimizing the sum of squared differences remains an underdetermined problem if we treat each value of the rate function as an independent unknown. However, under the assumption that biochemical rate functions are sufficiently smooth the problem can be solved by limiting the variability of the rate functions across neighbouring states (Materials & Methods).

To demonstrate how rate functions can be successfully inferred from partial observations of some components within a larger network we consider four different example networks that exhibited oscillations, bistability, fluctuation control, and noise enhancing feedback. Such markedly different global dynamics was already achievable with non-linear three-component feedback networks while conserving the reaction rates for one of the components, see Fig. 2 (left panels). We thus present the performance of our algorithm when applied to simulation data from those simple systems. But the algorithm can equally be applied to much larger systems with hundreds of variables as long as the local interactions are qualitatively known.

**FIG. 2.**
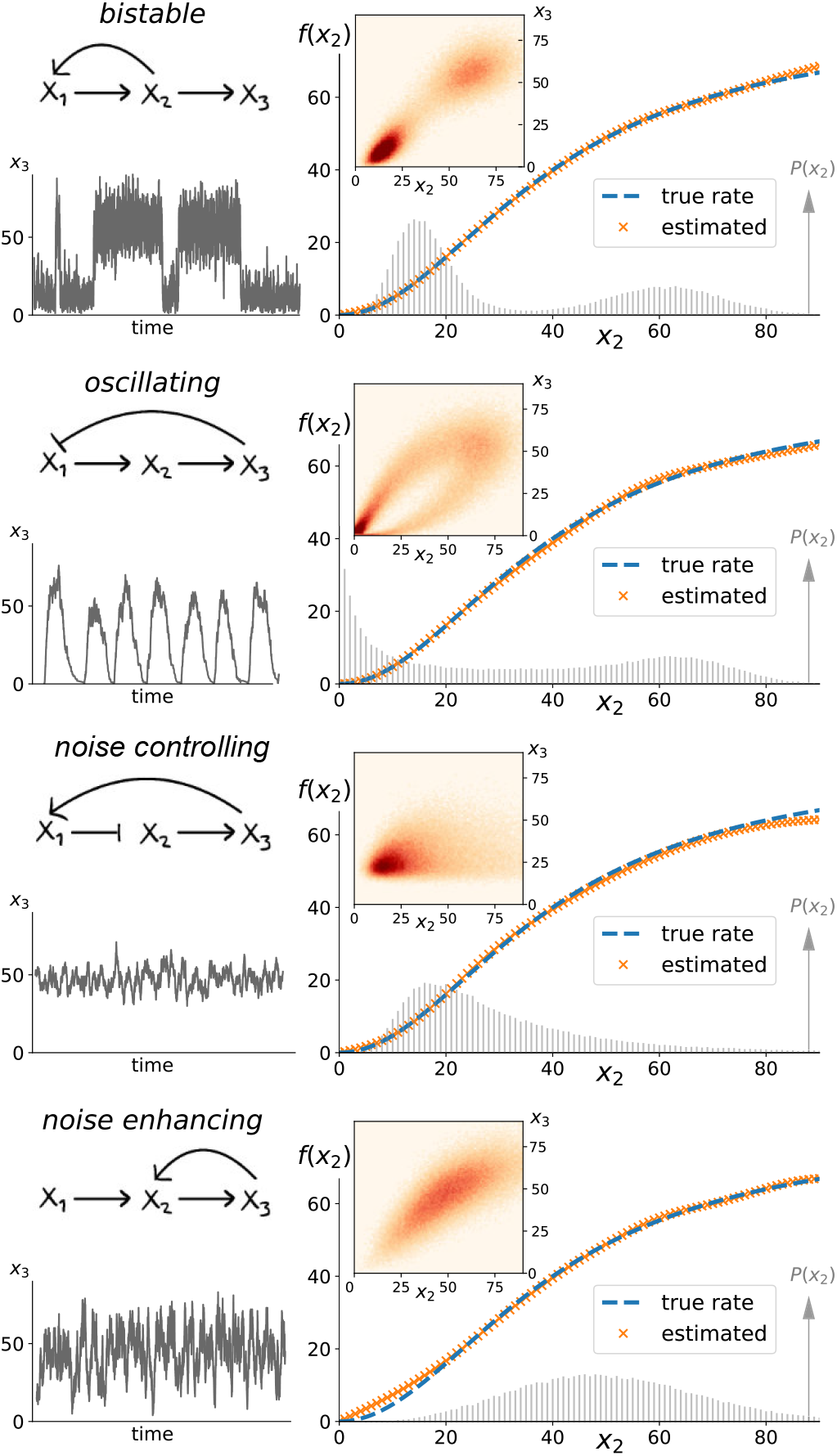
Probability flux balances can determine biochemical rates regardless of global network dynamics. Fixing how the production rate *f*(*x*_2_) of X_3_ depends on X_2_-levels, we considered four different global network topologies within the class defined by Eq. (3), that exhibit diverse system dynamics and variability in X_3_ (left column). The insets of the right column depict numerically observed joint distributions *P*(*x*_2_, *x*_3_) corresponding to 100,000 independent snapshots. Although probability distributions differed greatly between the four systems, Eq. (4) could identify the functional dependence of the production rate of X_3_ based on the numerical convex optimization algorithm detailed in the Materials & Methods. We find near perfect agreement between the inferred rate (orange crosses) and the true rate function (dashed blue line) regardless of a system’s global dynamics. This inference of *f*(*x*_2_) does not utilize any temporal information, its only input is the stationary joint probability distribution between the two components of interest. It relies on observing fluctuations across a wide range of X_2_-states as illustrated by the shaded probability distribution *P*(*x*_2_) with deviations occurring where X_2_ was rarely or never observed. While the degradation rate of X_3_ was assumed to be known, no information about how its production rate depends on X_2_, or the dynamics of X_1_, X_2_ was used.

### Numerical proof-of-principle examples

The conserved part of our test systems that we want to reconstruct from simulation data corresponds to a stereotypical biochemical reaction rate in cells. In particular, we specified that the production of component X_3_ is affected by X_2_ through a Hill-type function *f*(*x*_2_) and X_3_ molecules are degraded independently leading to the following class of reaction systems

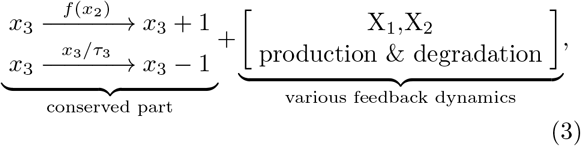

where *τ*_3_ denotes the average life-time of component X_3_ and 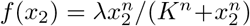. For test examples presented in Fig. 2 (dashed blue lines in right panels) we chose *τ*_3_ = 1 and λ = 80, *n* = 2, *K* = 40.

The reaction dynamics of the other variables, i.e., how X_1_, X_2_ affect each other and how they are affected by X_3_ were chosen to achieve diverse system dynamics and are specified in the Materials & Methods. Regardless of the X_1_, X_2_-dynamics, the probability balance equation Eq. (2) applied to the specified reaction in Eq. (3) imply that the above systems must satisfy the following probability balance relations at stationarity

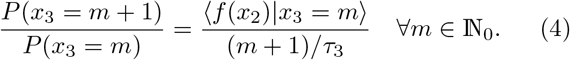

Although Eq. (4) is reminiscent of detailed balance, individual backwards and forward reaction fluxes do not need to balance in general for systems that do not operate at thermodynamic equilibrium such as cellular processes. The condition we exploit here is that the marginal probability distribution does not change at stationarity, and thus that each state must on average balance incoming and outgoing probability fluxes. The contrast with detailed balance is directly apparent in systems with dimeric degradation of X_3_ as discussed in a later section.

As detailed in the Materials & Methods, we employed a numerical algorithm to approximately solve Eq. (4) for *f*(*x*_2_) and thus infer how the X_3_ production rate depends on X_2_ from the observed *P*(*x*_2_, *x*_3_). To do so we generated exact realizations of the above stochastic processes using the standard Doob-Gillespie algorithm [18, 19]. We then sampled X_2_, X_3_ from the numerically observed stationary distribution to generate an observed joint probability from *N* = 100,000 independent samples. Any information about the dynamics of X_1_ was discarded because our method does not utilize any information beyond the pairs of components under consideration.

Applying a convex optimization algorithm to Eq. (4) using the numerically observed *P*(*x*_2_, x_3_) led to near perfect inference for the production rate of X3 in all systems as illustrated by the orange crosses in Fig. 2. These numerical proof-of-concept examples thus illustrate how the balance equations Eq. (4) can be used to reconstruct the functional form of *f*(*x*_2_) from pairwise observation of X_2_, X_3_ in the absence of any temporal information and independent of any information about the vastly different global system dynamics.

### Sampling requirements

Information cannot be created from nothing and the above inference cannot determine rates for states that were never observed. In practice, making additional assumptions to fill in gaps, such as monotonicity or the functional form of *f*(*x*_2_), could prove useful (and are easily incorporated into the algorithm), but here we want to illustrate the core of the inference quality based solely on the convex optimization of Eq. (4). We thus define an error heuristic *E* to quantify the quality of our inference by weighting errors in the rate function by the probability of the system to have been observed in that state, relative to the overall average of the rate function 〈*f*_true_〉:

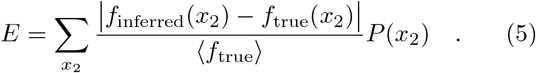

To illustrate how the relative time-scale of X_2_ and X_3_ affect this inference error E we consider the above “noise enhancing” system (Materials & Methods) for which changing lifetimes did not introduce different system dynamics. For such systems, inferring *f*(*x*_2_) is straight-forward when the variability of X_2_ is slow such that X_3_ has enough time to adjust to X_2_-levels and the conditional average 〈*x*_3_|*x*_2_〉 directly identifies the production rate of X_3_ (SI). For faster upstream fluctuations the effect of X_2_-variability on X_3_ decreases and the inference of the production rate becomes more challenging. However, compared to the naive statistical approach of interpreting conditional averages as rates, our inference algorithm based on Eq. (4) reliably identifies the correct rate function even when the time-scale of X_2_-fluctuations is fast relative to X_3_, see Fig. 3B. When the upstream variably becomes more than an order of magnitude faster than X3 our inferred rate function deviates significantly from the true one when inferred from a joint probability distribution constructed from *N* = 100, 000 samples. However, even in this unfavourable regime with a 40-fold separation of time-scale between the upstream and downstream variable such that the conditional average 〈*x*_3_|*x*_2_〉 levels-off, *N* = 5 × 10^6^ were enough sample observations to correctly infer *f*(*x*_2_), see Fig. 3D.

**FIG. 3.**
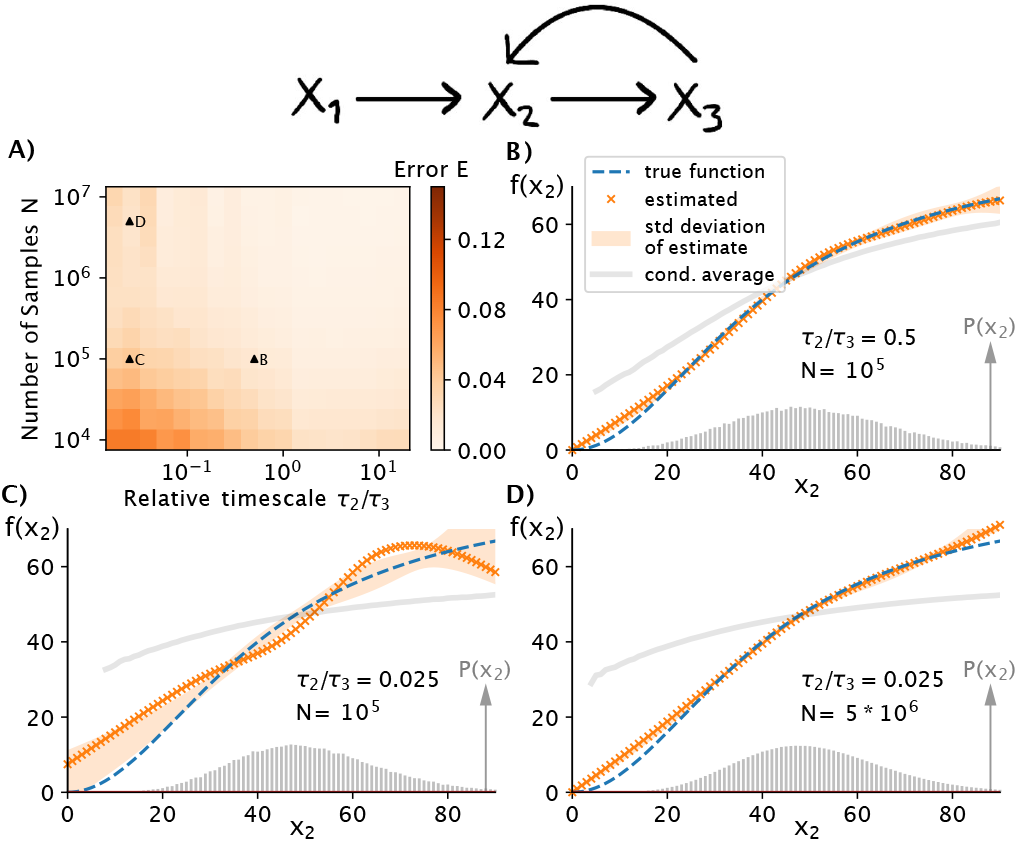
Experimentally achievable sampling leads to accurate inference even when fast upstream variability masks rate dependencies. A) The number of data points required to successfully infer *f*(*x*_2_) from an empirical *P*(*x*_2_, *x*_3_) depends on the relative time-scales between the two components of interest. Simulations of the noise enhancing three-component system (Materials & Methods) show that several thousand measurement samples can be enough to reliably infer the rate dependence *f*(*x*_2_) when upstream fluc-tuations are relatively slow, i.e., *τ*_2_ > *τ*_3_. B) When upstream fluctuations are fast, the downstream variable does not have time to adjust and the conditional average 〈*x*_3_|*x*_2_〉 no longer follows *f*(*x*_2_) as indicated by the grey line. In contrast, our inference method based on Eq. (4) accurately estimates the actual rate function from *N* = 100, 000 samples. The orange crosses depict an example of one inference while the shaded area displays the standard deviation of individual inferences from different samples of the same process. C) As the upstream fluctuations in X_2_ become faster, the inference gets worse when using the same number of sampling points. However, even for systems in which X_2_, and X_3_ time-scales are separated 40-fold, the production rate *f*(*x*_2_) can be accurately inferred from *N* = 5 × 10^6^ samples (panel D). Such sampling is experimentally achievable using flow-cytometry approaches to characterize single cell heterogeneity [20].

### Upregulated *vs*. downregulated production rates

While the above examples exhibit vastly different global system dynamics the functional form of the production rate of X_3_ was conserved across all systems. Next, we demonstrate that our inference method works for arbitrary Hill-type functions for the production rate. We simulated the noise enhancing three-component system (Materials & Methods) from Fig. 3 with differently shaped Hill-functions 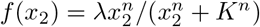 by systematically varying the parameters *K, n*. The tested range of parameters reflects biologically relevant different shapes, including negative *n* corresponding to X_2_ suppressing X_3_, small values of *n* such that X_3_ is barely affected by X_2_, as well as strongly cooperative effects with *n* → ±4. As illustrated in Fig. 4A, the inference works satisfactorily for *N* = 100, 000 across a broad range Hill-functions with specific examples of successful inference depicted in Fig. 4C,D,E. Note, that those rate functions cannot be quantified for states that were never (or extremely rarely) observed.

**FIG. 4.**
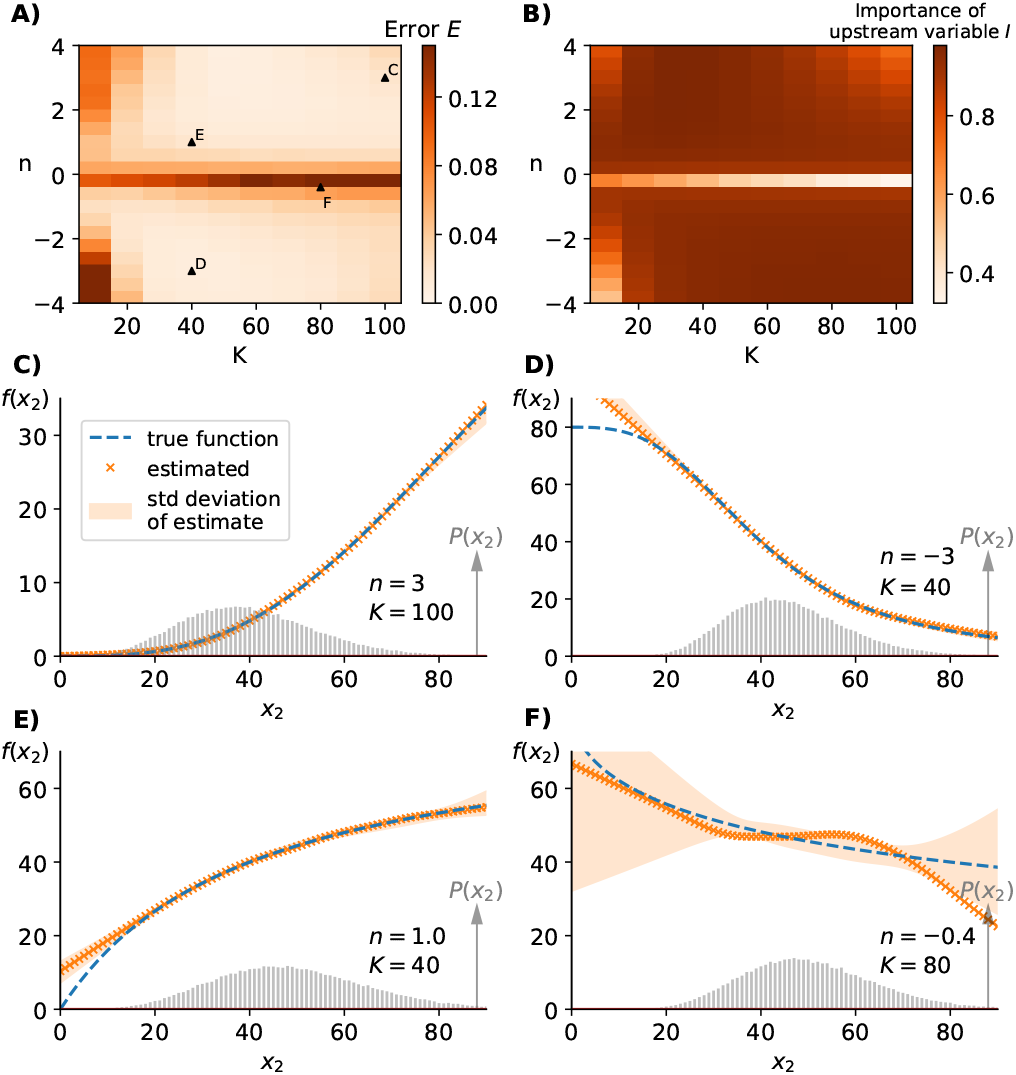
Different shapes of rate functions can be inferred. A) Changing the shape of the production rate 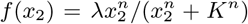 by varying *K* and *n* while keeping time-scales fixed and equal, we find that the inferred reaction rates using Eq. (4) and *P*(*x*_2_, *x*_3_) agree well with the true rate for *N* = 100, 000 samples across a broad range of the parameter regime. Data shown are for the same noise enhancing three-component system (Materials & Methods) as in Fig. 3. B) Unsatisfactory inference of *f*(*x*_2_) corresponds to regimes in which the upstream variable has only little influence on the downstream fluctuations as quantified by the relative importance term *I* defined in Eq. (6). C,D,E) Successful inference examples for different Hill-functions including repressing effects of X_2_ on X_3_. Any deviations from the true rate function are in the region in which the system is rarely or never observed. F) Example of unsatisfactory inference when the reaction rate varies only little over the majority of observed states resulting in a small effect of X_2_-fluctuations on X_3_-levels. The poor inference is also highlighted by the extremely broad shaded region indicating the standard deviation of inferred *f*(*x*_2_) for identical systems subject to different random sampling.

To determine the cause of unsatisfactory inferences as illustrated by an example in Fig. 4F, we utilize the general noise propagation relation [16] to describe all systems within the class of Eq. (3)

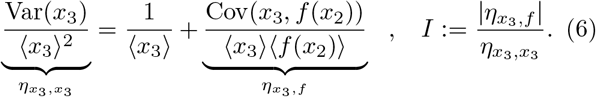

Here, *I* quantifies how much of an effect X_2_-fluctuations have on X_3_-variability. We find, that regions of unsatisfactory inference correspond to systems in which the upstream variability has only a small effect on X_3_ (compare panels A and B of Fig. 4) Inference in the regime where *n* ≈ 0 can be significantly improved through a simple cross-validation step as discussed in a later section and illustrated in Fig. 7B.

### Non-linear degradation rates

In all of the above systems, X_3_-molecules were degraded in a first-order reaction as is commonly the case for cellular components [21, 22] and would be approximately true for all cellular components that are not actively degraded but effectively diluted by cellular growth [23]. Next, we demonstrate that our inference method works equally well for systems in which X_3_ undergoes non-linear degradation reactions.

Analogous to Fig. 4, we varied the shape of the production rate *f*(*x*_2_) in a class of noise enhancing (Materials & Methods) systems with the following conserved part

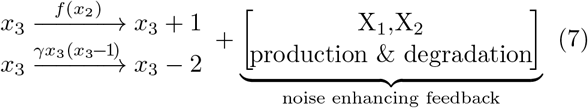

where a non-linear degradation rate corresponding to a dimerization event was added. We again find that our inference method works reliable for most parameters, see Fig. 5A, with unsatisfactory results corresponding again to parameter regimes in which the upstream variable has only a marginal effect on the downstream fluctuations.

**FIG. 5.**
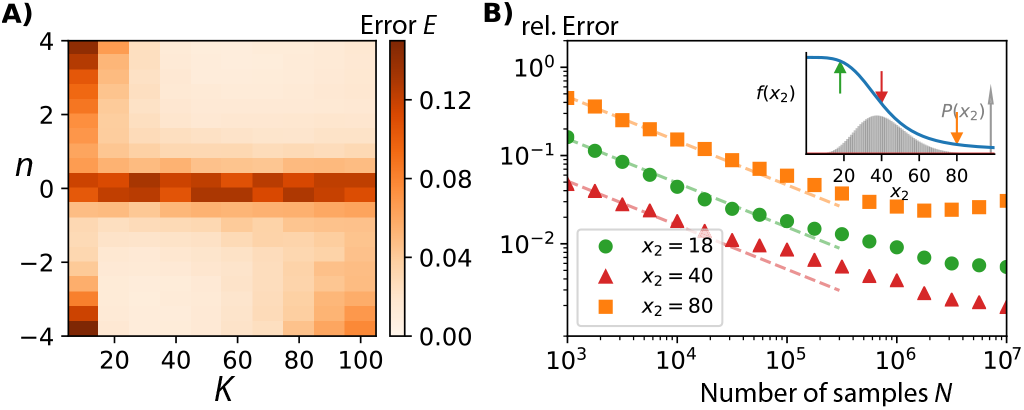
Non-linear degradation does not affect the inference. A) For simulated systems with non-linear degradation of X_3_-molecules as defined in Eq. (7), the inference quality is satisfactory when using empirically determined joint probability distributions *P*(*x*_2_, *x*_3_) from *N* = 100,000 samples. Plotted are the inference error *E* for different production rates 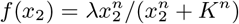. Poor inference corresponds to parameter regimes in which the upstream variable X_2_ has only a negligible effect on the downstream variable X_3_, i.e., when *K* or *n* are small. Data are for the same noise enhancing three-component system as in Fig. 3 with the only difference that the degradation of X3 is now non-linear. B) For a given state, the relative error of the inferred reaction rate *f*(*x*_2_) initially decreases 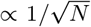 (dashed lines) as the number of sampling points *N* increases. However, for large *N*, the relative error levels off, with higher probability states reaching a lower plateau than those only visited rarely. The inset depicts the true function and the specific states considered here, as well as the resulting probability distribution of *x*_2_.

Additionally, we analyzed how the inference quality of *f*(x_2_) behaves for individual states. As intuitively expected, we find that the inference error initially decreases 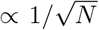 as the number of samples *N* increases. However, for large *N* a plateau becomes apparent that is most severe for the most rarely observed states, Fig. 5B. This behaviour was generally observed across different classes of systems (SI). Explicitly accounting for the effects of sampling error may lead to lower plateaus for the estimation errors with more advanced statistical methods to invert Eq. (4) but are beyond the scope of the current work.

Note, in these systems the component of interest X_3_ degrades as a dimer, such that its specified degradation rate in Eq. (7) is non-linear, and the reaction eliminates two molecules at a time. The probability balancing Eq. (2) therefore no longer involves just neighbouring states and detailed balance is broken. Instead the above systems must satisfy the following balance equations

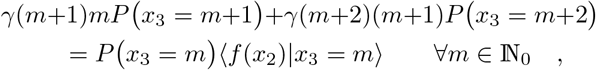

that were used in the inference algorithm.

### Experimental Measurement Noise

Due to finite sampling and unavoidable measurement noise, empirically observed probability distributions will not perfectly reproduce the stationary distributions of the underlying chemical reaction network. Next, we analyze how measurement noise affects our inference method by explicitly accounting for small absolute and relative error terms as well as systemic undercounting of molecules.

To simulate “empirically observed” probability distributions of the above noise enhancing system (Materials & Methods) we resampled from the exact stationary distribution *P*(*x*_2_, *x*_3_) while adding a two-dimensional normally distributed error with zero mean and a standard deviation of *σ*_abs_ = 1, 3, 8 molecules respectively (SI). For small absolute errors, we find that the inference method still succeeds to satisfactorily determine the original rate function, see Fig. 6B.

**FIG. 6.**
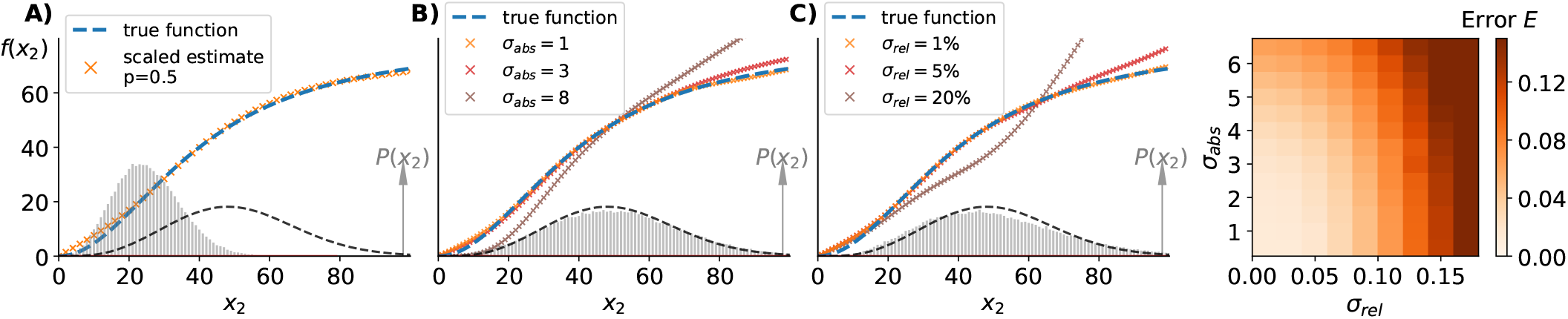
Small measurement noise does not prohibit accurate inference. A) Simulated “empirical” data (histogram) with binomial undercounting in which each molecule is detected with a fixed probability *p*. Our inference algorithm identifies the correct reaction rate if we simply multiply the measured molecule numbers by 1/*p* again to obtain the input function (dashed blue line). If we do not know *p* then the algorithm identifies the correct shape *f*(*x*_2_) but cannot identify the correct scale of X_2_ over which it varies. B) Simulated “empirical” data (grey histogram) with added absolute measurement errors modelled as a two-dimensional Gaussian with zero mean and standard deviation of *σ_abs_* = 8. This distribution is not identical to the theoretical one (dashed black line) and leads to significant deviations in the inference (brown crosses). For smaller absolute errors we observe satisfactory inference, as illustrated by the data for *σ_abs_* = 1, 3. C) Simulating relative measurement errors by multiplying “observed” samples from the exact stationary distribution with a random number from a two-dimensional Gaussian with mean one and standard deviation of *σ_rel_* = 0.01, 0.1, 0.2. For relative errors less than 10% we found satisfactory inference but larger errors led to unacceptable estimates for the production rate *f*(*x*_2_) because of the significant deviation of the “empirical” distribution (grey histogram) from the exact stationary distribution (dashed black line) illustrated for *σ_rel_* =0.2. Explicit de-convolution steps [24] for known types of measurement noise may significantly improve the inference performance of future algorithms based on Eq. (2). D) Quantifying the inference error for absolute and relative measurement errors. Relative measurement errors larger than 10% led to unsatisfactory inference.

Furthermore, we analyzed the effect of relative measurement noise by multiplying each sampled data point with a two-dimensional normally distributed error term to simulate a relative error of 1%, 5%, 20% in the observed variables respectively (SI). In its current form, our inference is significantly affected by large multiplicative noise because it causes the “measured” probability distribution to differ significantly from the underlying stationary distribution of the stochastic process, see Fig. 6C. Future variants of our inference algorithm may potentially improve on this by performing explicit de-convolution steps to estimate stationary state distributions from experimentally recorded ones before exploiting Eq. (2). In fact, measuring error due to probabilistic undercounting can be exactly accounted for by determining the probability *p* to detect a specific molecule, and applying our inference method to the re-scaled probability distribution, see Fig. 6A.

### Weakly connected components

Our presented method relies on knowing that one component directly affects the production rate of another. We thus obtained unsatisfactory inferences when the upstream variable barely affects the downstream variable, as illustrated in the regime when *n* → 0 or *K* ≪ 〈*x*_2_〉, see Fig. 4A.

Breakdown of satisfactory inference in that regime can be prevented by explicitly considering the possibility that *f*(*x*_2_) is approximately constant across the observed stochastic fluctuations of X_2_. Following a standard crossvalidation approach, we can use one half of the observed data as a “training set” [25]. Using this subset of data we infer *f*(*x*_2_) using our usual unconstrained method but additionally perform a constrained optimization for constant production rates, i.e., we find the best *f*(*x*_2_) = λ for some λ > 0. This by itself will not pick a constant production rate over the freely optimized *f*(*x*_2_) because the latter has many more degrees of freedom. However, because the free optimization overfits sampling errors when minimizing deviations of Eq. (4) in the regime in which *f*(*x*_2_) is approximately constant, it will do relatively worse than a constant production rate when applied to the “validation set” of the data. Incorporating this cross-validation approach into our inference methods as detailed in the Materials & Methods, removes the most unsatisfactory regime while leaving the successful inferences unaffected as illustrated in Fig. 7A (compare to Fig. 4A) with an example detection of constant *f*(*x*_2_) illustrated in Fig. 7B.

**FIG. 7.**
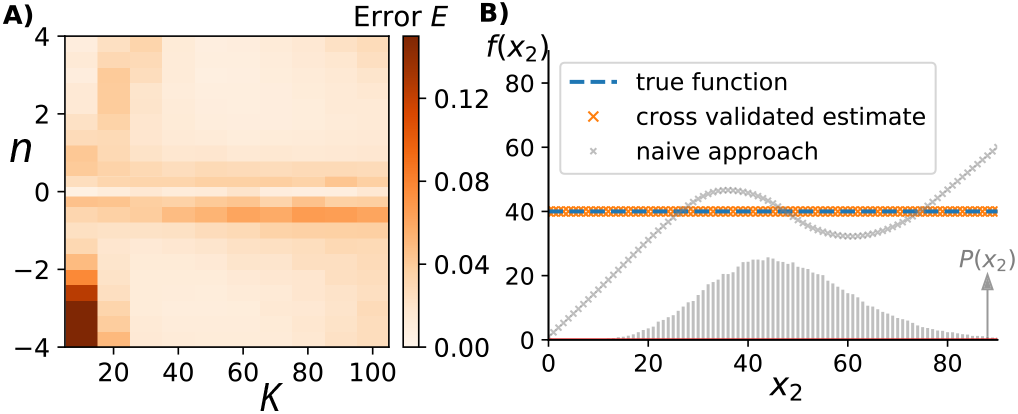
Cross-validation improves inference in regions with constant to near constant rates. A) Analogous to Fig. 4 we change the shape of the production rate 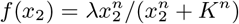 by varying *K* and *n*. Shown here is the error of the inferred *f*(*x*_2_) when applying our method with an additional cross-validation step as detailed in the Materials & Methods. The quality of the inference is significantly improved in the regime in which the production rate is essentially independent of the upstream variable around *n* ≈ 0. B) Directly comparing the cross-validated and freely inferred *f*(*x*_2_) when applied to a system in which the true production rate was constant. The cross-validated answer picks out the constant production rate, whereas the naive approach gives a varying estimate due to overfitting of Eq. (4).

Identifying constant production rates is a special case of the general problem of identifying which component affects which in complex biochemical reaction networks [26]. Future variants of such a cross-validation approach might thus prove useful in identifying the topology of network interactions based on Eq. (4).

## Discussion

Early aeronautical engineering faced a challenge analogous to that of current synthetic biology. While the qualitative requirements for motored flight were well known, designing a reliable flyer required a breakthrough in *quantitatively* understanding each subproblem through painstaking experimental measurements [27].

Similarly, designing reliable synthetic circuits in biology requires a quantitative description of biochemical reaction dynamics rather than qualitative network interaction models. Here, we present a method that promises to iteratively turn a qualitative model of biochemical interactions into a network of quantitative reaction rates. Given one arrow within a network of interacting components our method identifies the functional dependence of the actual reaction rate, one interaction at a time, from fluctuations of a subset of components without having to perturb the system.

Existing mechanistic modelling approaches often rely on temporal data [28–35] and generally have to be identified from data in one fell swoop [36–41] with all the reliability issues that come with optimizing over many degrees of freedom at once. While approaches that combine mechanistic models with Bayesian inference [42–45] can effectively account for significant noise in experimental data, they too rely on inferring parameters for fully defined mechanistic models all at once. Statistical approaches that rely on perturbations [7–11] can be straightforwardly applied to static snapshots of incompletely observed complex systems but fail to account for the dynamic ways in which one component affects another, and thus do generally not quantitatively describe physical interactions between components.

Our results show that we can reliably identify reaction rates independent of the larger network structures, in contrast to previous approaches in which small models of gene expression were inverted under the assumption they were correct as a whole and only a handful of parameter values needed to be determined [46–48]. The presented numerical proof-of-principle results establish that our presented algorithm works across a broad range of tested systems and that the number of required observations for algorithmic success is comparable to those accessible by modern experimental single-cell techniques in biology such as flow-cytometry that routinely measure millions of isogenic cells at a time.

## Materials & Methods

### Basic algorithm

To utilize probability distributions that contain sampling errors we write the balance equations (4) for our example systems as G**f** = **h**, where *G_ij_* = *P*(*x*_2_ = *j*, *x*_3_ = *i*), *f_i_* = *f*(*x*_2_ = *i*) and *h_i_* = (*i* + 1)*P*(*x*_3_ = *i* + 1)/*τ*_3_. To avoid overfitting the solution **f** to sampling noise we add a regularization term that effectively penalizes the discrete “second derivatives” *f*_*i*+2_ – 2*f*_*i*+1_ + *f_i_*. The effect of this regularization is to smoothen the resulting reaction rate function. Motivated by complex cellular processes where *f*(*x*_2_) represent biochemical reaction rates, we furthermore constrain the solution vector **f** to be non-negative. We thus solve the following optimization problem

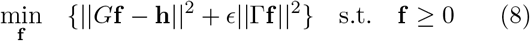

where Γ_*ij*_ = *δ_i, j_* – 2*δ*_*i, j*+1_ + *δ*_*i, j*+2_ is the regularization matrix and e corresponds to the strength of the smoothening. The strength of this regularization parameter affects the quality of the inference. We found 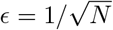 to lead to satisfactory results because it appropriately decreases in strength as the number of sampling points *N* increases. This regularization was used throughout the paper. In other applications, alternative heuristics to choose e may prove useful to achieve satisfactory results (SI). Note, that if the lifetime *τ*_3_ is unknown, the inference method will still correctly identify *τ*_3_ · *f*(*x*_2_) meaning that we get the correct shape of the reaction rate function up to an unknown scale-factor.

We solved Eq. (8) using a standard convex optimization approach which is guaranteed to converge to the optimal solution as a linear program [49]. Note, that it is straightforward to add further constraints, such as monotonicity, about the solution function *f*(*x*_2_). However, the results presented here do not make use of any additional assumptions beyond Eq. (8).

### Cross-validation algorithm

In order to avoid overfitting systems when production rates are approximately constant we compared our inferred production rate against a constant one as follows: We divided the sampling data into two equally sized sets, and applied our inference method first to the “training” set and then checked against a constant production rate using the second “validation” set [25]. To compare the two rates we calculated the violation of Eq. (4) as the sum of all squared errors. If the constant production rate’s error was smaller or within a 5% margin of the freely fitted production rate from the training set, we determined a constant production rate to be the best inference and optimized it over the whole data. Otherwise we applied our regular inference method to the full data set instead.

### Definition of example systems

We considered simple three-component systems in which the production and degradation rates of X_1_, X_2_, X_2_ took the following form

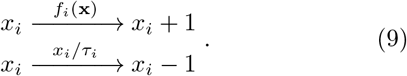

In particular, we simulated the following example systems depicted in Fig. 2 where the production and degradation rate for X_3_ was kept the same as specified in the main text while the other components were subject to the following dynamics.

#### Bistable system

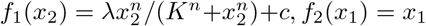 with λ = 50, *n* = 6, *K* = 37, *c* = 15 and life-times *τ*_1_ = *τ*_2_ = 1.

#### Oscillating system

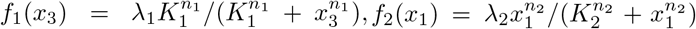 with λ_1_ = 50000, *n*_1_ = 10, *K*_1_ = 0.1, λ_2_ = 80, *n*_2_ = 1, *K*_2_ = 100 and life-times *τ*_1_ = *τ*_2_ = 1.

#### Noise controlling system

*f*_1_(*x*_3_) = λ_1_*x*_3_, 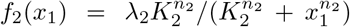 with λ_1_ = 50, λ_2_ = 3000, *n*_2_ = 10, *K*_2_ = 10 and life-times *τ*_1_ = 50, *τ*_2_ = 1.

#### Noise enhancing system

*f*_1_ = 5, 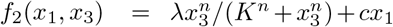 with λ = 25, *n* = 4, *K* = 50, *c* =8 and life-times *τ*_1_ = *τ*_2_ = 1.

Data in Figs. 3-7 correspond to the above noise enhancing system with modifications as specified in the main text. The systems with non-linear degradation rate *γx*_3_(*x*_3_ – 1) were simulated with *γ* = 2.

## Acknowledgements

We thank M. Assaf for the suggestion of the regularization term and N. Lord and C. Zechner for valuable feedback on the manuscript. We thank Raymond Fan, Brayden Kell, Seshu Iyengar, Euan Joly-Smith for many helpful discussions and suggestions to improve the inference method. This work was supported by the Natural Sciences and Engineering Research Council of Canada and a New Researcher Award from the University of Toronto Connaught Fund. NL was supported by research awards from the Israel Council for Higher Education (VATAT) and a Ben-Gurion University Postdoctoral Fellowship. TW gratefully acknowledges financial support through a Deutschlandstipendium.

## Author contributions

T.W., N.L. derived the results and all authors wrote the manuscript.

## Correspondence

Correspondence and requests should be addressed to A.H.

