## Supplemental Information for "Quantifying biochemical reaction rates from static population variability within complex networks"

#### This PDF file includes:

Supplementary text
Figs. S1 to S3
SI References

### Supporting Information Text

### 1. Simulation details

All simulations were performed using the standard Doob-Gillespie (1, 2) algorithm until each reaction occurred at least $10^8$ times. The resulting high-fidelity distributions from these simulations were considered as the underlying 'true' distribution for each respective system. To obtain the distributions used to test our inference algorithm, we randomly sampled $N$ individual data points ($N = 100,000$ if not otherwise specified) from the 'true' distributions corresponding to $N$ independent snapshots of a given system. For this sampling we used the *numpy.random.choice* function in Python.

**Reaction dynamics for numerical proof-of-principle examples in Fig. 2 of main text.** Below we specify the parameters used to simulate the numerical proof-of-principle examples introduced in Figure 2 of the main text. These parameters do not necessarily correspond to realistic cellular networks but were chosen to achieve a distinct set of system dynamics to make the mathematical point that we can successfully infer $f(x_2)$ for diverse system dynamics across comparable range for $X_2$-variability. All variables are subject to the following reactions

$$\begin{aligned} x_i &\xrightarrow{f_i(\mathbf{x})} x_i + 1 \\ x_i &\xrightarrow{x_i/\tau_i} x_i - 1 \end{aligned} \qquad [S.1]$$

with a linear degradation rate dependent on the lifetime $\tau_i$ of a specific molecule $X_i$ and a production rate $f_i(\mathbf{x})$ dependent on the state vector $\mathbf{x}$.

As mentioned in the main text the production rate of $X_3$was kept the same for all systems: $f_3(x_2) = \lambda_3\, x_2^{n_3}/(K_3^{n_3} + x_2^{n_3})$ with a lifetime of $\tau_3 = 1$ and the Hill function parameters $\lambda_3 = 80, n_3 = 2, K_3 = 40$. The other rate functions were chosen as follows:

- Bistable system

$$\begin{aligned} f_1(x_2) =&\ \lambda_1 \frac{x_2^{n_1}}{K_1^{n_1} + x_2^{n_1}} + c_1\ ; && \lambda_1 = 50,\ n_1 = 6,\ K_1 = 37,\ c_1 = 15 \\ f_2(x_1) =&\ \lambda_2 x_1\ ; && \lambda_2 = 1 \\ \text{Lifetimes:} & && \tau_1 = \tau_2 = \tau_3 = 1 \end{aligned}$$

- Oscillating system

$$\begin{aligned} f_1(x_3) =&\ \lambda_1 \frac{K_1^{n_1}}{K_1^{n_1} + x_3^{n_1}}\ ; && \lambda_1 = 50000,\ n_1 = 10,\ K_1 = 0.1 \\ f_2(x_1) =&\ \lambda_2 \frac{x_1^{n_2}}{K_2^{n_2} + x_1^{n_2}}\ ; && \lambda_2 = 80,\ n_2 = 1,\ K_2 = 100 \\ \text{Lifetimes:} & && \tau_1 = \tau_2 = \tau_3 = 1 \end{aligned}$$

- Noise controlling system

$$\begin{aligned} f_1(x_3) =&\ \lambda_1 x_3\ ; && \lambda_1 = 50 \\ f_2(x_1) =&\ \lambda_2 \frac{K_2^{n_2}}{K_2^{n_2} + x_1^{n_2}}\ ; && \lambda_2 = 3000,\ n_2 = 10,\ K_2 = 10 \\ \text{Lifetimes:} & && \tau_1 = 50,\ \tau_2 = \tau_3 = 1 \end{aligned}$$

- Noise enhancing system

$$f_1 = \lambda_1 \; ; \qquad \lambda_1 = 5$$

$$f_2(x_1, x_3) = \lambda_2 \frac{x_3^{n_2}}{K_2^{n_2} + x_3^{n_2}} + c_2 x_1 \; ; \qquad \lambda_2 = 25,\; n_2 = 4,\; K_2 = 50,\; c_2 = 8$$

$$\text{Lifetimes:} \qquad \tau_1 = \tau_2 = \tau_3 = 1$$

**Reaction dynamics for example systems in Fig. 3-5 of main text.** To illustrate how the inference method depends on time-scales and sampling (Fig. 3 of the main text) we simulated the "noise enhancing system" as defined above while logarithmically varying the life-time $\tau_3 = 1.5^k$ with $k$ ranging from $-7$ to 10. In order to keep system averages $\langle x_3 \rangle$ comparable while varying the time-scales we simultaneously adjusted the value of $\lambda_3$ such that $\lambda_3 \tau_3 = 80$ remained constant.

To illustrate the dependence of the inference method on reaction rate function $f(x_2)$ (Fig. 4 of the main text) we simulated the above "noise enhancing system", with $f_3(x_2) = \lambda_3 \, x_2^{n_3}/(K_3^{n_3} + x_2^{n_3})$ for varying $K_3, n_3$ and fixed $\lambda_3 = 80, \tau_3 = 1$.

To illustrate that the nature of the degradation rate does not significantly affect the inference (Fig. 5 of the main text) we simulated the above "noise enhancing system" but replaced the degradation reaction of $X_3$ with

$$x_3 \xrightarrow{\gamma x_3 (x_3 - 1)} x_3 - 2 \qquad \text{[S.2]}$$

with a fixed value of $\gamma = 2$.

### 2. Modeling measurement errors – Fig. 6 of the main text

In the main text we analyzed how measurement noise affects our inference method by accounting for absolute and relative error terms as well as undercounting of molecules. The following error models $(x_2, x_3) \mapsto (x_2', x_3')$ were applied to each sampled data point:

- abolute error

$$(x_2', x_3') = (\mathcal{N}(x_2, \sigma_{abs}), \mathcal{N}(x_3, \sigma_{abs})),$$

- relative error

$$(x_2', x_3') = (x_2 \cdot \mathcal{N}(1, \sigma_{rel}), x_3 \cdot \mathcal{N}(1, \sigma_{rel})),$$

- binominal undercounting

$$(x_2', x_3') = (\mathcal{B}(x_2, p), \mathcal{B}(x_3, p)),$$

where $\mathcal{N}(\mu, \sigma)$ refers to a normal distribution with a mean of $\mu$ and standard variation of $\sigma$. $\mathcal{B}(n, p)$ refers to a binomial distribution where $p$ corresponds to the probability that a specific molecule gets detected. All resulting $(x_2', x_3')$ pairs were rounded to integers and pairs with negative molecule numbers were ignored.

When applying our inference method to simulated data with binomial undercounting the resulting rate function depends on the undercounted variable $x_2'$. The correct rate function was obtained by rescaling the inferred rate function argument with the detection probability $p$ such that $f(x_2) = f(x_2'/p)$.

### 3. Derivation of probability balance relations for specific systems

Our starting point is the following probability balance equation which must hold for any given molecular component $X_i$ of a complex network that reaches a stationary state in which its probability distribution $P_{ss}(x_i)$ no longer changes. This equations follows trivially from the master equation and was explicitly discussed previously elsewhere (3).

$$0 = \sum_k \Big[\langle r_k(\mathbf{x})|x_i = m - d_{ki}\rangle P_{ss}(x_i = m - d_{ki}) - \langle r_k(\mathbf{x})|x_i = m\rangle P_{ss}(x_i = m)\Big] \quad \forall m \in \mathbb{N}_0. \qquad \text{[S.3]}$$

**General systems with linear degradation of $X_3$.** First we derive the invariant balance equation for the following class of systems introduced in the main text.

$$\underbrace{\begin{array}{ccc} x_3 & \xrightarrow{f(x_2)} & x_3+1 \\ x_3 & \xrightarrow{x_3/\tau_3} & x_3-1 \end{array}}_{\text{specified part}} + \underbrace{\left[\begin{array}{c} X_1,X_2 \\ \text{production \& degradation} \end{array}\right]}_{\text{various feedback dynamics}}. \qquad \text{[S.4]}$$

Applying Eq. S.3 to the specific dynamics of Eq. S.4 leads to

$$\langle f(x_2)|x_3 = m-1\rangle P(x_3 = m-1) + \frac{m+1}{\tau_3}P(x_3 = m+1) = \langle f(x_2)|x_3 = m\rangle P(x_3 = m) + \frac{m}{\tau_3}P(x_3 = m) \quad \forall m \in \mathbb{N}_0. \qquad \text{[S.5]}$$

where we have dropped the subscript to denote the stationary state probability distribution for notational convenience. For biochemical reaction systems, for the lowest state $m = 0$ the relationship simplifies to

$$\langle f(x_2)|x_3 = 0\rangle P(x_3 = 0) = \frac{1}{\tau_3}P(x_3 = 1), \qquad \text{[S.6]}$$

because molecular numbers cannot be negative (which implies that $P(x_3 = -1) = 0$). Combining Eq. S.5 with Eq. S.6 then gives rise to the following recursion relation

$$\langle f(x_2)|x_3 = m\rangle P(x_3 = m) = \frac{m+1}{\tau_3}P(x_3 = m+1) \quad \forall m \in \mathbb{N}_0, \qquad \text{[S.7]}$$

which is Eq. (4) presented in the Main Text.

To get the matrix form of these relations which we use in our optimization algorithm, we re-wrote the conditional averages in Eq. S.7

$$\sum_i f(x_2 = i)P(x_2 = i, x_3 = m) = \frac{m+1}{\tau_3}P(x_3 = m+1) \quad \forall m \in \mathbb{N}_0, \qquad \text{[S.8]}$$

where $P(x_2, x_3)$ is the joint probability of $X_2$ and $X_3$.

In experiments and computational simulations there will always be a maximum value $m_{max}$ for which $P(x_3 = m) = 0 \quad \forall m > m_{max}$ such that Eq. S.8 can be written in matrix form

$$G\mathbf{f} = \mathbf{h} \qquad \text{[S.9]}$$

where $G_{ij} = P(x_2 = j, x_3 = i)$, $f_i = f(x_2 = i)$ and $h_i = (i+1)P(x_3 = i+1)/\tau_3$.

**General systems with non-linear degradation of $X_3$.** The derivation of the probability flux balance relations for systems in which $X_3$ is degraded as a dimer

$$\underbrace{\begin{array}{ccc} x_3 & \xrightarrow{f(x_2)} & x_3+1 \\ x_3 & \xrightarrow{\gamma x_3(x_3-1)} & x_3-2 \end{array}}_{\text{specified part}} + \underbrace{\left[\begin{array}{c} X_1,X_2 \\ \text{production \& degradation} \end{array}\right]}_{\text{irrelevant feedback dynamics}} \qquad \text{[S.10]}$$

is equivalent to the one in the previous section. First, we apply Eq. S.3 the specified dynamics of Eq. S.10 to obtain

$$\begin{aligned}&\langle f(x_2)|x_3=m-1\rangle P(x_3=m-1)+\gamma m(m-1)P(x_3=m)\\=&\langle f(x_2)|x_3=m\rangle P(x_3=m)+\gamma(m+2)(m+1)P(x_3=m+2)\quad\forall m\in\mathbb{N}_0.\end{aligned}\tag{S.11}$$

This is a second order recurrence equation that simplifies in the states $m=0$ and $m=1$ for which the degradation rate is zero, such that for $m=0$

$$\langle f(x_2)|x_3=0\rangle P(x_3=0)=2\gamma P(x_3=2),\tag{S.12}$$

because $P(x_3=-1)=0$. For $m=1$ we obtain

$$\begin{aligned}&\langle f(x_2)|x_3=0\rangle P(x_3=0)-\langle f(x_2)|x_3=1\rangle P(x_3=1)+6\gamma P(x_3=3)\\&=-\langle f(x_2)|x_3=1\rangle P(x_3=1)+2\gamma P(x_3=2)+6\gamma P(x_3=3)=0.\end{aligned}\tag{S.13}$$

Combining these, then leads to the following general balance relation

$$\langle f(x_2)|x_3=m\rangle P(x_3=m)=\gamma m(m+1)P(x_3=m+1)+\gamma(m+1)(m+2)P(x_3=m+2)\quad\forall m\in\mathbb{N}_0,\tag{S.14}$$

as stated in the main text.

### 4. Heuristic approach to determine the regularization parameter

Following the assumption that biochemical reaction rates are reasonably smooth we introduced a regularization term in our optimization in order to penalize large jumps in the reaction rate $f(x_2)$

$$\min_{\mathbf{f}}\ \{\underbrace{||G\mathbf{f}-\mathbf{h}||^2}_{\text{unbiased objective function}}+\underbrace{\epsilon||\Gamma\mathbf{f}||^2\}}_{\text{regularization term}}\quad\text{s.t.}\quad\mathbf{f}\geq 0,\tag{S.15}$$

where the regularization term with $\Gamma_{ij}=\delta_{i,j}-2\delta_{i,j+1}+\delta_{i,j+2}$ essentially corresponds the discrete second derivative of $f(x_2)$. Throughout the paper we used $\epsilon=1/\sqrt{N}$ which appropriately decreases the importance of the regularization term as the data become more precise. However, it is possible that for other systems another value for the regularization parameter may be advantageous. For completeness, we describe a general method to determine an "optimal" value for $\epsilon$ for arbitrary cases.

The goal is to pick a regularization parameter which is strong enough to smoothen our estimated reaction rate without introducing too strong a linear bias to the optimization. This can be achieved by determining the value of the unbiased objective function of Eq. S.15 after optimization as a function of the regularization parameter. In Fig. S1A we illustrate the result for the "bistable system" introduced earlier. We see that after a certain value for $\epsilon$ there is a sharp increase of the objective function which means that the optimization gets heavily biased by the regularization term beyond a certain value. As a heuristic we thus propose to pick a value for the regularization parameter just before the sharp increase. Large values of $\epsilon$ essentially force the resulting $f(x_2)$ to be a straight line leading to the plateau in the objective function.

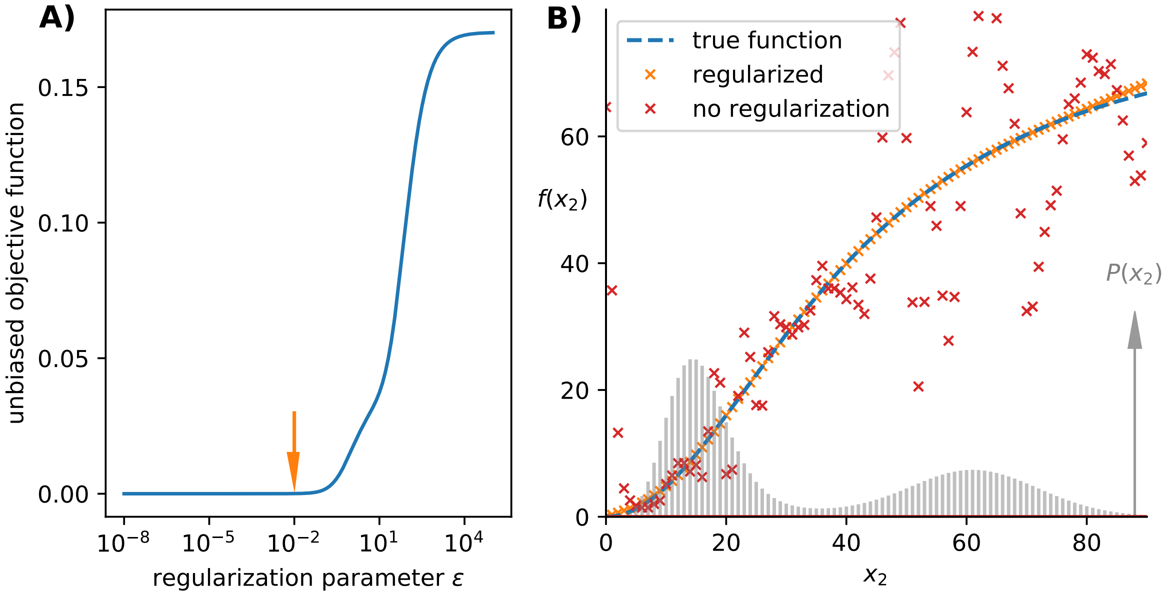


**Fig. S1. The optimal value for the regularization parameter can be heuristically determined through its effect on the objective function.** A) Choosing a regularization parameter strong enough to smoothen the estimated reaction rate without introducing a significant bias leads to the best inference. In this case it was determined to be $\epsilon = 0.01$ (orange arrow). B) The regularization is essential to achieve optimal results. Shown here are the inferred rate with the chosen $\epsilon = 0.01$ (orange crosses) and without regularization, i.e., $\epsilon = 0$ (red crosses).

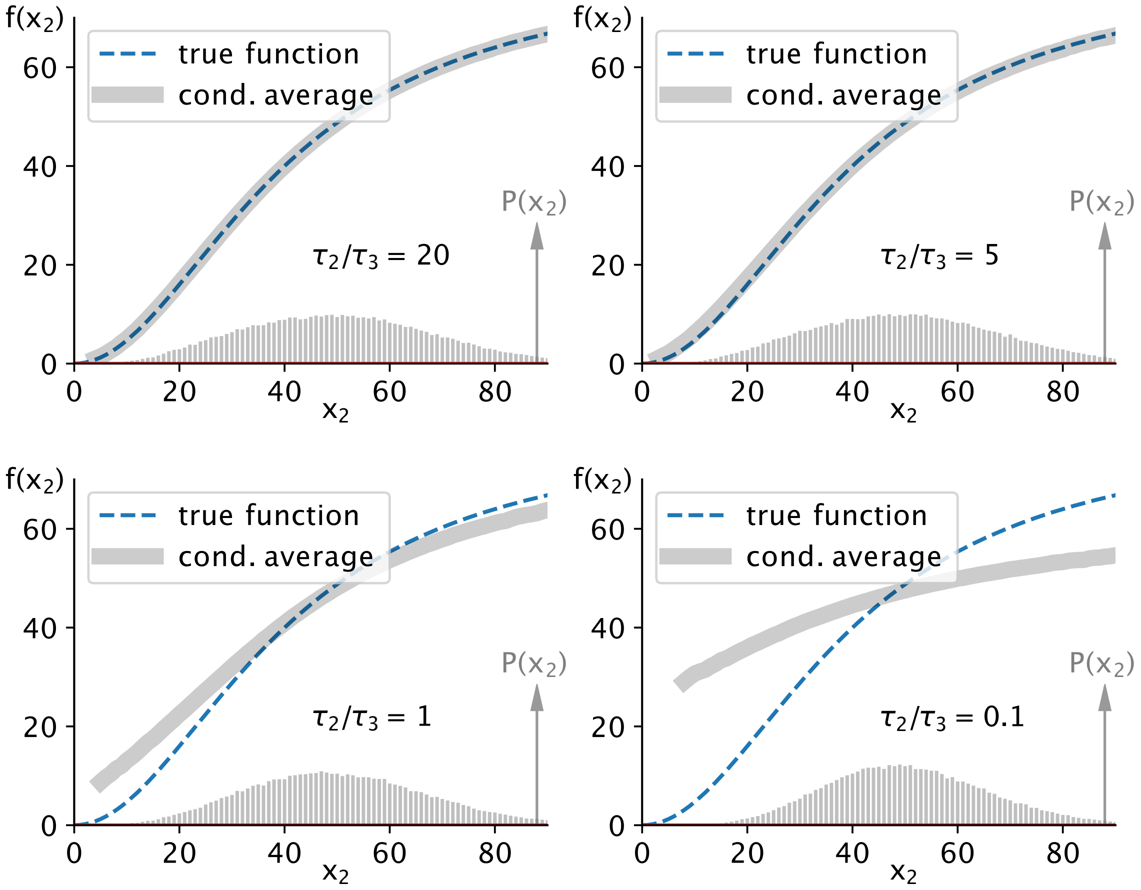


**Fig. S2. For fast upstream variables the conditional average follows the reaction rate function.** We consider the "noise enhancing" system defined above with changing lifetimes ratios. For such systems, inferring $f_3(x_2)$ is straightforward when $\tau_2 \gg \tau_3$ and the variability of $X_2$ is slow enough for $X_3$ to adjust to $X_2$-levels and the conditional average $\langle x_3|x_2\rangle$ directly identifies the production rate of $X_3$. In contrast, when upstream variability in $X_2$ is fast compared to the life-time of $X_3$ the conditional average no longer follows the production rate.

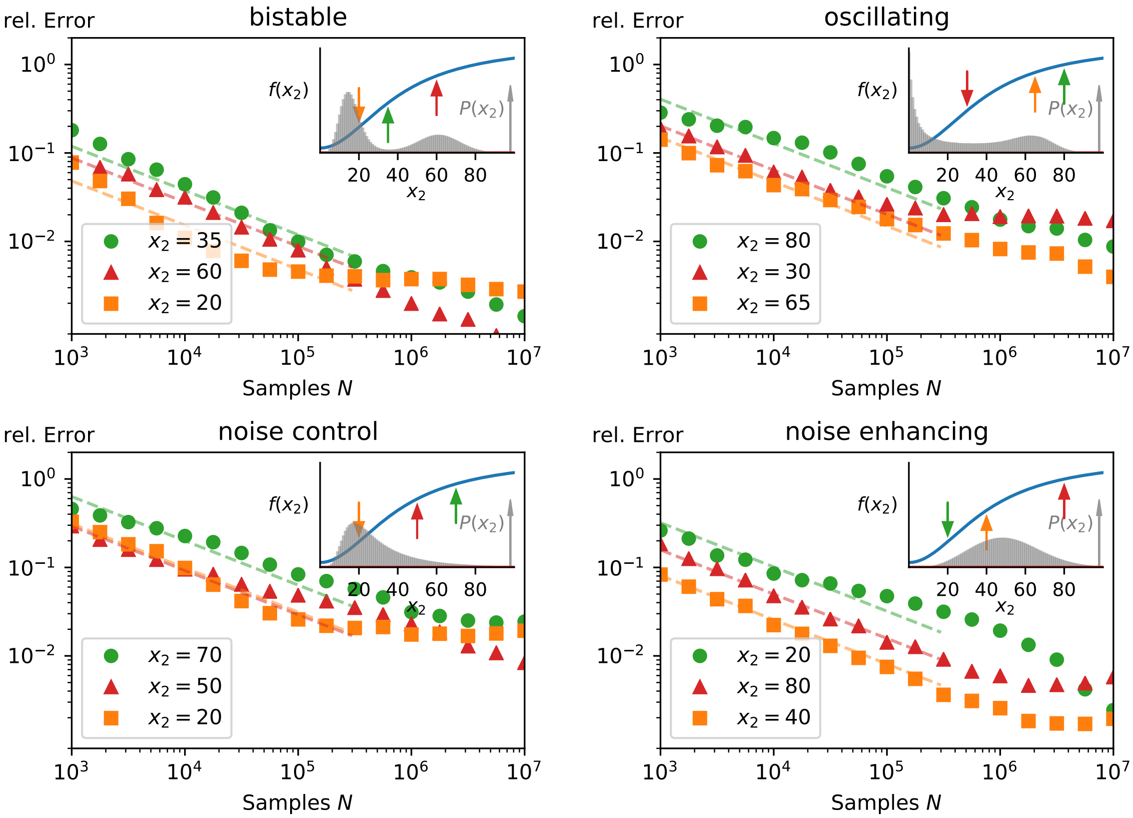


**Fig. S3. Inference quality behaviour for individual states.** For each of the four systems of Fig. 2 in the main text we analyzed the error behaviour for a given state $x_2 = m$. Initially, the relative error of the inferred reaction rate $f(x_2)$ approximately decreases $\propto 1/\sqrt{N}$ (dashed lines) as the number of sampling points $N$ increases. For large $N$, the relative error levels off, with higher probability states reaching a lower plateau than those only visited rarely. For very large sampling some of the errors can be seen to increase slightly as our choice of $\epsilon = 1/\sqrt{N}$ becomes suboptimal. The inset depicts the true function, the specific states considered here, and the respective probability distribution of $X_2$.
